# Stimulus-dependent orientation strategies in monarch butterflies

**DOI:** 10.1101/2021.10.14.464409

**Authors:** Myriam Franzke, Christian Kraus, Maria Gayler, David Dreyer, Keram Pfeiffer, Basil el Jundi

**Affiliations:** University of Wuerzburg, Biocenter, Zoology II, Würzburg, Germany; Lund University, Department of Biology, Lund Vision Group, Lund, Sweden

**Keywords:** insect, vision, landmark, lepidoptera, stripe fixation, attraction behavior, compass orientation

## Abstract

Insects are well-known for their ability to keep track of their heading direction based on a combination of skylight cues and visual landmarks. This allows them to navigate back to their nest, disperse throughout unfamiliar environments, as well as migrate over large distances between their breeding and non-breeding habitats. The monarch butterfly (*Danaus plexippus*) for instance is known for its annual southward migration from North America to certain trees in Central Mexico. To maintain a constant flight route, these butterflies use a time-compensated sun compass for orientation which is processed in a region in the brain, termed the central complex. However, to successfully complete their journey, the butterflies’ brain must generate a multitude of orientation strategies, allowing them to dynamically switch from sun-compass orientation to a tactic behavior toward a certain target. To study if monarch butterflies exhibit different orientation modes and if they can switch between them, we observed the orientation behavior of tethered flying butterflies in a flight simulator while presenting different visual cues to them. We found that the butterflies’ behavior depended on the presented visual stimulus. Thus, while a dark stripe was used for flight stabilization, a bright stripe was fixated by the butterflies in their frontal visual field. If we replaced a bright stripe by a simulated sun stimulus, the butterflies switched their orientation behavior and exhibited compass orientation. Taken together, our data show that monarch butterflies rely on and switch between different orientation modes, allowing them to adjust orientation to the actual behavioral demands of the animal.

## Introduction

Orientation in space is an essential ability for animals to find food, escape from predators, or return to their nest. To achieve this, insects exhibit a number of different orientation mechanisms, ranging from the simple straight-line orientation of dung beetles (Dacke et al., 2021; el Jundi et al., 2019) to more complex behaviors such as path integration of ants and bees (Collett and Collett, 2000; Heinze et al., 2018) or long-distance migration of lepidopterans (Grob et al., 2021; Hu et al., 2021; Merlin and Liedvogel, 2019; Warrant et al., 2016). One striking example of a migrating insect is the monarch butterfly (*Danaus plexippus*) (Reppert and de Roode, 2018; Reppert et al., 2016). Each fall millions of these butterflies migrate from the northern USA and Canada over more than 4,000 km to their overwintering habitat in Central Mexico. To keep a constant direction over this enormous distance, these animals rely on the sun for orientation (Froy et al., 2003; Heinze and Reppert, 2011; Mouritsen and Frost, 2002; Reppert, 2006). In combination with time-of-day information from circadian clocks in the brain (Sauman et al., 2005) and/or the antennae (Guerra et al., 2012; Merlin et al., 2009) this allows the butterflies to maintain a directed course throughout the day. Beside the sun, additional cues, such as the celestial polarization pattern (Reppert et al., 2004) or the Earth’s magnetic field (Guerra et al., 2014; Wan et al., 2021) seem to play a role during the migration but their relevance for the butterfly’s compass is still not fully understood (Stalleicken et al., 2005).

As in other insects, the central complex of monarch butterflies serves as an internal compass during spatial orientation (el Jundi et al., 2014; Heinze and Reppert, 2011; Heinze et al., 2013; Pfeiffer and Homberg, 2014). Compass neurons in this brain region are sensitive to multiple simulated skylight cues (Heinze and Reppert, 2011; Nguyen et al., 2021) and encode the animal’s heading with respect to a sun stimulus (Beetz et al., 2021). As shown previously, a sun stimulus – represented by a green light spot – can be employed in behavioral laboratory experiments in monarch butterflies (Franzke et al., 2020). Similar experiments in the fruit fly *Drosophila melanogaster* demonstrated that these insects exhibit a menotactic behavior with respect to a simulated sun. This means that the fruit fly maintains any arbitrary heading relative to the sun (Giraldo et al., 2018). Interestingly, closed-loop experiments showed that as soon as the activity of central-complex neurons was genetically deactivated, the flies kept the simulated sun in their frontal visual field, resembling vertical stripe fixation behavior (Giraldo et al., 2018). This attraction behavior does not depend on whether the flies are confronted with a bright stripe on a dark background or the inverted visual scene (Maimon et al., 2008). Although the biological function of the fly’s attraction behavior is not fully understood, it is speculated that the flies interpret this cue as a landing or feeding site (Maimon et al., 2008). Whether monarch butterflies adjust their orientation strategy depending on the visual stimulus is not known. However, to successfully display a large repertoire of behaviors, the orientation network in the butterfly’s brain needs to possess the capacity to flexibly switch between different orientation circuitries that may operate in parallel in the brain. This would, for instance, allow a flying butterfly to change from compass orientation based on skylight cues to attraction based on a visual landmark or an odor plume similar to what has been found for homing desert ants (Buehlmann et al., 2013).

To study the monarch butterflies’ behavioral repertoire, we recorded the orientation behavior of flying butterflies, tethered at the center of an LED-flight simulator, while we provided different visual cues (dark stripe, bright stripe, and sun stimulus) to the animals. We found that the butterflies used the dark stripe for flight stabilization based on optic-flow information. A bright stripe on the other hand evoked a simple attraction behavior towards the stimulus. In contrast, a simulated sun was used by the butterflies to maintain a constant angle with respect to the stimulus. We furthermore found that the butterflies switched between compass orientation and attraction behavior during flight. Taken together, our results show that monarch butterflies display different orientation modes that allow them to dynamically switch between different behaviors while navigating through their environment.

## Material and Methods

### Experimental animals

Pupae of the monarch butterfly (*Danaus plexippus*) were ordered from Costa Rica Entomology Supply (butterflyfarm.co.cr) and kept in an incubator (HPP 110 and HPP 749, Memmert GmbH+Co. KG, Schwabach, Germany) at 25°C and 80% relative humidity and under a 12:12 h light:dark cycle. After the animals eclosed, the adult butterflies were transferred to a flight cage inside a separate incubator (I-30VL, Percival Scientific, Perry, IA, USA) with a 12 h:12 h light:dark cycle. While the relative humidity was constant at about 50%, the temperature was set to 25°C during light phases and 23°C during dark phases. Feeders inside the flight cage were filled with 15% sucrose solution and provided *ad libitum* food to the butterflies.

### Preparation

We used female and male adult butterflies and prepared them in the morning prior to each experiment. The scales of the butterflies’ thorax were removed and a tungsten stalk (0.508×152.4 mm, Science Products GmbH, Hofheim, Germany) was glued (multi-purpose impact instant contact adhesive, EVO-STIK, Bostik Ltd, Stafford, UK) dorsally to it. After preparation, the animals were individually kept in clear plastic containers with access to 15% sucrose solution and transferred to a dark chamber for at least three hours.

### Flight simulator

We used a flight simulator similar to the ones described previously (Dreyer et al., 2018a; Dreyer et al., 2018b; Dreyer et al., 2021). To record the heading directions of individual butterflies, the tungsten wire on the animals’ thorax was connected to an optical encoder (E4T miniature Optical Kit Encoder, US Digital, Vancouver, WA, USA). This allowed the butterflies to rotate at the center of the flight simulator and freely choose any heading. Butterflies that stopped flying for more than four times during an experiment were excluded from the study. The heading direction of the animals was recorded with an angular resolution of 3 deg and a temporal resolution of 200 ms using a data acquisition device (USB4 Encoder Data Acquisition USB Device, US Digital, Vancouver, WA, USA) and a computer with the corresponding software (USB1, USB4: US Digital, Vancouver, WA, USA). To present visual stimuli to the butterflies, the inner surface of the flight simulator was equipped with a circular array of 2048 RGB LEDs (128*16 APA102C LED Matrix, iPixel LED Light Co.,Ltd, Baoan Shenzhen, China) or green high power LEDs (LZ1-00G102, OSRAM, San Jose, CA, USA). A custom written python script was used to control the color and intensity of all LEDs of the LED arena via a raspberry pi (Raspberry Pi 3 Model B, Raspberry Pi Foundation, UK).

### Stimuli

In all experiments we presented the butterflies with one or multiple different visual stimuli. To produce them, the intensity and color of each LED of the arena was adjusted as summarized in the following table:

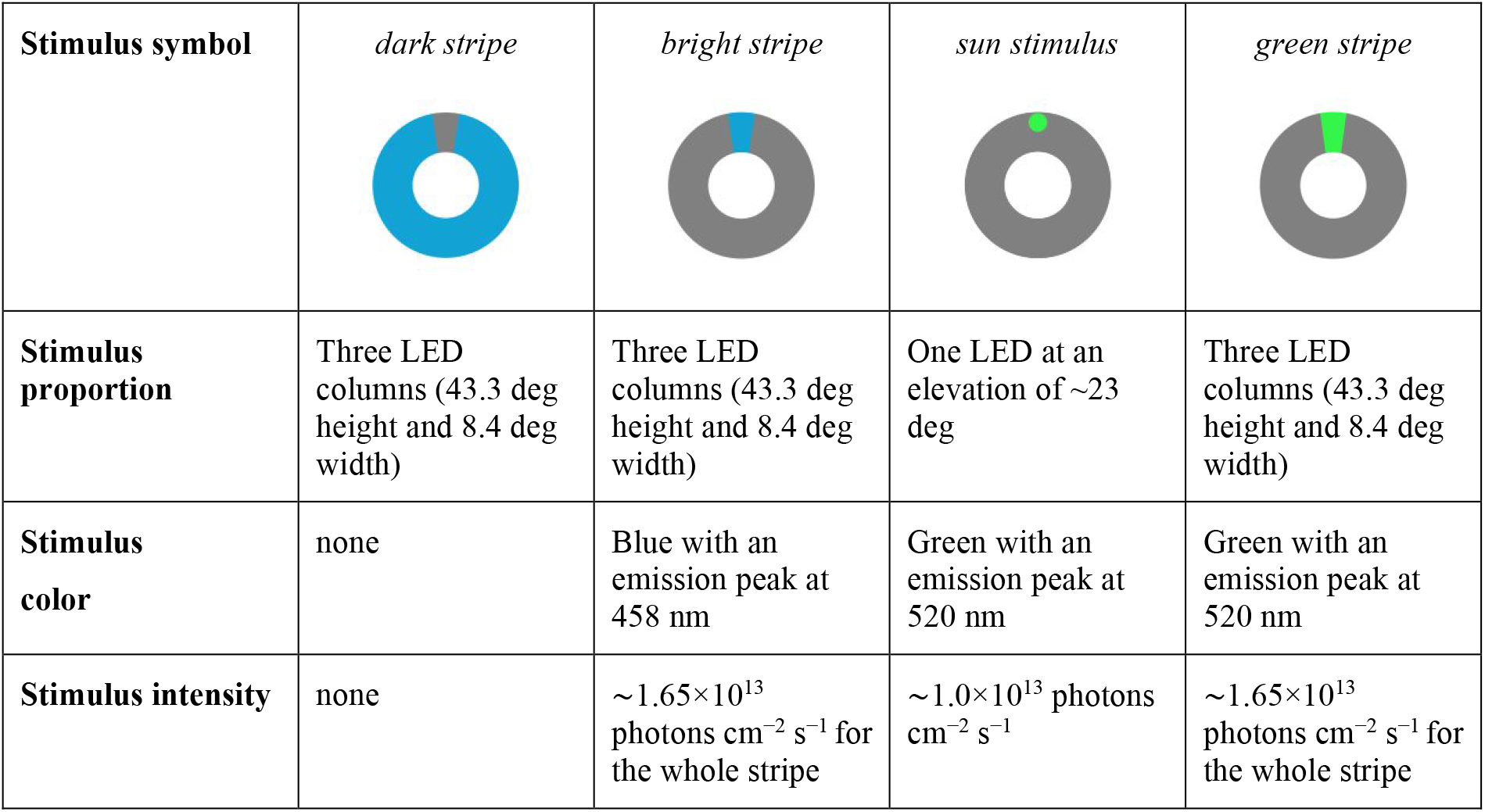
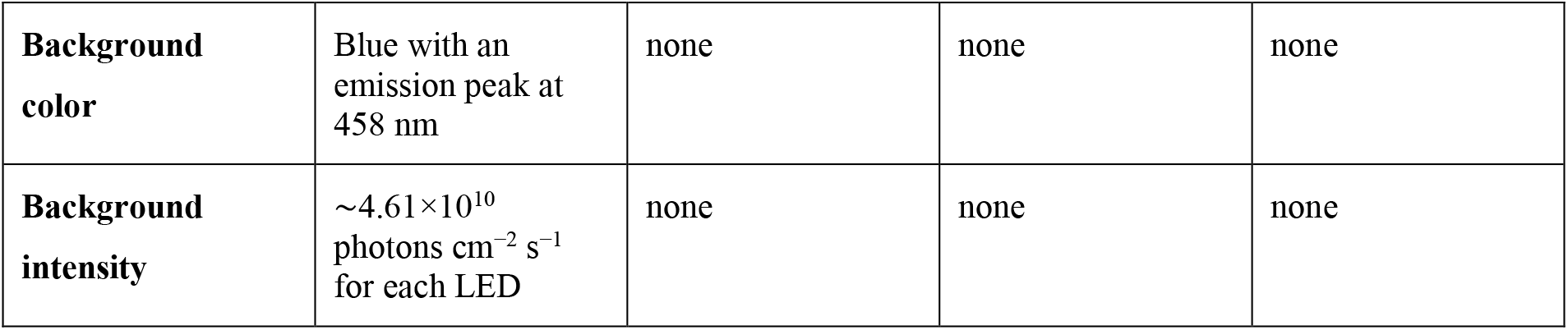

| Stimulus symbol | *dark stripe* | *bright stripe* | *sun stimulus* | *green stripe* |
| --- | --- | --- | --- | --- |
| Stimulus proportion | Three LED columns (43.3 deg height and 8.4 deg width) | Three LED columns (43.3 deg height and 8.4 deg width) | One LED at an elevation of ~23 deg | Three LED columns (43.3 deg height and 8.4 deg width) |
| Stimulus color | none | Blue with an emission peak at 458 nm | Green with an emission peak at 520 nm | Green with an emission peak at 520 nm |
| Stimulus intensity | none | $\sim 1.65 \times 10^{13}$ photons $\text{cm}^{-2} \text{s}^{-1}$ for the whole stripe | $\sim 1.0 \times 10^{13}$ photons $\text{cm}^{-2} \text{s}^{-1}$ | $\sim 1.65 \times 10^{13}$ photons $\text{cm}^{-2} \text{s}^{-1}$ for the whole stripe |
| **Background color** | Blue with an emission peak at 458 nm | none | none | none |
| **Background intensity** | $\sim 4.61 \times 10^{10}$ photons $\text{cm}^{-2} \text{s}^{-1}$ for each LED | none | none | none |

### Orientation with respect to one cue

In the first set of experiments, we presented one cue (stripe or simulated sun) as an orientation reference to the butterflies. In all experiments, the heading direction of a single animal was recorded over eight minutes. To ensure that the butterflies used the displayed cue for orientation, we turned the visual scenery by 180 deg every two minutes and studied if the animals followed the relocation of the cue. We alternated the start position of the stimulus between 0 deg and 180 deg, starting at 0 deg for half of the animals and at 180 deg for the other half.

First, we investigated whether monarch butterflies can use a landmark for orientation by setting all LEDs of the arena to blue while three LED columns were turned off which generated a dark stripe on a bright background (Fig. 1A; experiment: *dark stripe*). 28 butterflies were then individually connected via the tungsten wire to the encoder at the arena’s center and were allowed to orient by changing their heading direction with respect to the landmark. As a control, we also performed an experiment with 22 individuals, in which the animals did not perceive any visual cue for orientation. Therefore, all LEDs were set to blue (experiment: *no cue*; same data as in Franzke et al., 2020).

**Fig. 1.**
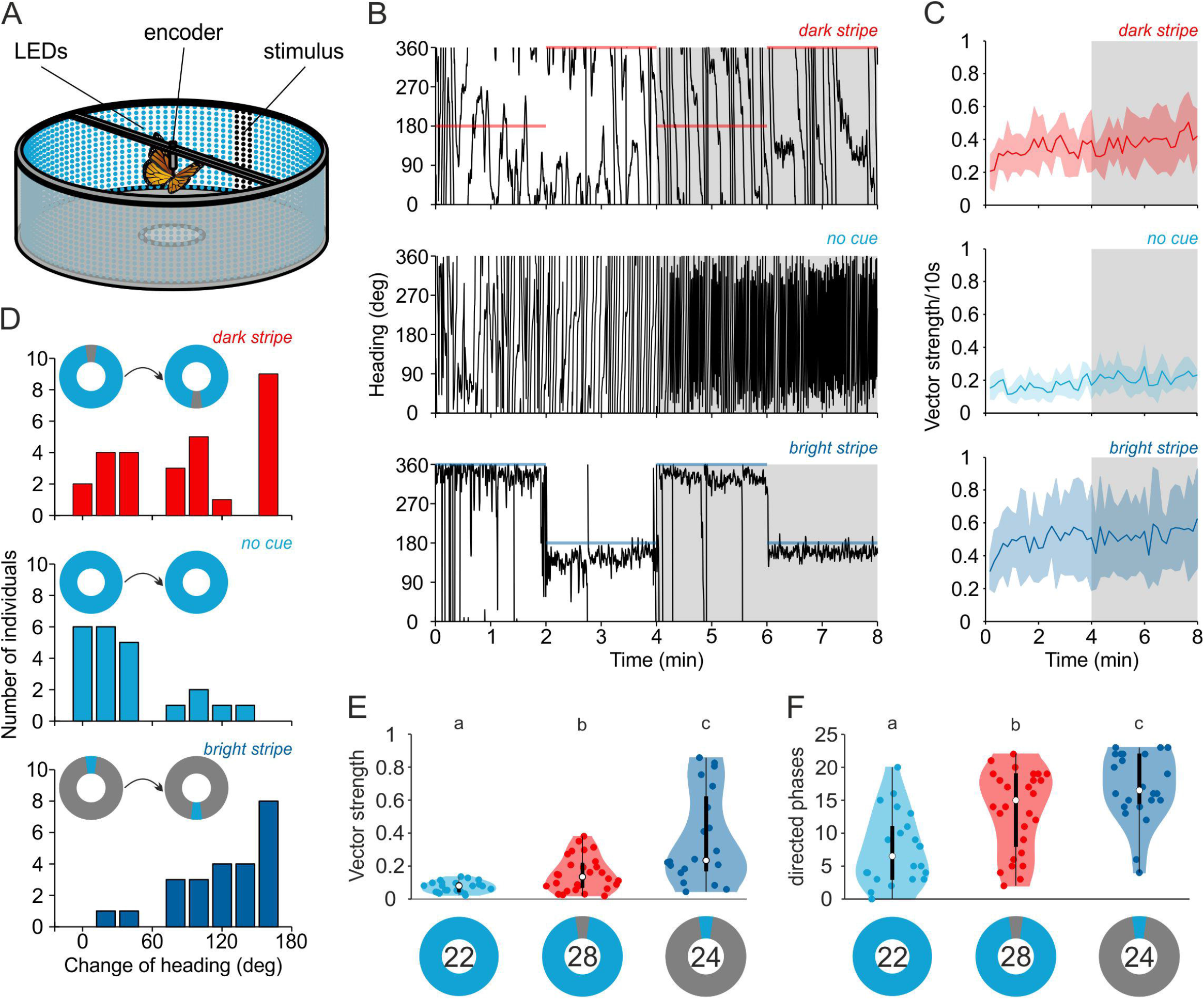
Landmark orientation to a vertical stripe. (A) Schematic illustration of a monarch butterfly tethered at the center of a flight simulator equipped with 2048 RGB-LEDs. While presenting visual stimuli to the butterflies, their heading directions were recorded using an optical encoder. (B) Flight trace of exemplary butterflies that were flying in the LED arena with respect to a dark stripe on a blue background (*dark stripe*), with all LEDs turned blue (*no cue*), or with a bright, blue stripe on a dark background as orientation reference (*bright stripe*). Colored, horizontal lines indicate the position of the vertical stripe at either 0 or 180 deg. The gray boxes indicate the 4-min sections that were used for further analysis. (C) The mean vector strength *r* (bin size: 10 s) over the entire eight-minute experiments shows that the butterflies’ orientation performance increased over time. The animals were better oriented when a dark stripe was added to a bright background (*dark stripe*, first panel; *no cue*, second panel) and performed the best when a bright stripe was presented (*bright stripe*, third panel). Shaded areas indicate the 25–75% quantile. The gray boxes indicate the 4-min sections that were used for further analysis. (D) Change of heading (bin size: 20 deg) between the last two phases of an experiment (indicated by the gray boxes in B) when the stimulus was relocated by 180 deg. Fewer animals changed their heading when no cue was available (*no cue*), but most animals followed a relocation of the stripes by more than 60 deg (*dark stripe, bright stripe*). (E) The vector strength (bin size: 4 min) was significantly higher when a dark stripe was added to a blue background (P=0.009, χ^2^= 6.76, Kruskal-Wallis test) but not as high as when a bright stripe was displayed (*dark stripe* against *bright stripe:* P=0.004, χ^2^=8.52, Kruskal-Wallis test). White dots indicate the median vector strength. The black boxes show the interquartile range and thin black lines extend to the 1.5 x interquartile range. Colored dots show the individual data points and shaded area represents their density. Letters indicate a significant difference between the tested groups. (F) Significantly more directed phases (vector strength >0.249; bin size: 10s) were observed when a dark stripe was added to the blue background (P= 0.001, χ^2^= 10.31, Kruskal-Wallis test). This number increased when a bright stripe (P=0.032, χ^2^=4.59, Kruskal-Wallis test) was presented. Plot conventions as in E.

Next, we inverted the visual scenery by turning all LEDs off with the exception of three LED columns which were set to blue. We recorded the headings of 24 butterflies presented with this bright stripe (experiment: *bright stripe*).

Finally, we investigated which orientation strategy monarch butterflies display when they were flying with respect to a simulated sun. Previous studies revealed that a green light cue is interpreted as the direction towards the sun by several insects (Edrich et al., 1979; el Jundi et al., 2015; Rossel and Wehner, 1984). Therefore, we presented 20 animals a simulated sun by turning one bright green LED on (experiment: *sun stimulus*). To test if the spectral content of our stimuli (green sun stimulus vs. blue stripe) had an impact on the orientation behavior of the butterflies, we repeated the experiments with the bright stripe with 20 butterflies. This time, the stripe changed its color every two minutes of flight from green to blue and *vice versa*. Again, the position and color of the stimulus was alternated between each butterfly. This means a quarter of the animals first experienced a green stripe at 0 deg (green stripe 0/blue stripe 180/green stripe 0/ blue stripe 180 deg) while a quarter of the butterflies started with a green stripe at 180 deg first (green stripe 180/ blue stripe 0/green stripe 180/blue stripe 0 deg). The remaining animals perceived the blue stripe at either 0 deg (blue stripe 0/green stripe 180/blue stripe 0/green stripe 180 deg) or 180 deg first (blue stripe 180/green stripe 0/blue stripe 180/green stripe 0 deg).

In a final step, we investigated if the butterflies changed their orientation behavior when we changed the stimulus during a butterfly’s directed flight. Therefore, we repeated the experiment with the blue vs. green stripe with 33 animals. However, instead of changing the color, this time we changed the appearance of the stimulus presenting a bright, green stripe for one minute followed by a sun stimulus for another one-minute phase (*or vice versa*). The animals’ orientation was recorded over two minutes and the position and stimulus order (sun stimulus to stripe, stripe to sun stimulus) was alternated for each butterfly.

### Orientation within an ambiguous scenery

In the second set of experiments, we presented two identical stimuli to the butterflies, that were set 180 deg apart from each other. In all experiments, the position of the stimuli was relocated, this time by 90 deg, every two minutes for a total of eight minutes. Again, we alternated the start position of the stimulus between 0/180 deg and 90/270 deg, starting at 0/180 deg for half of the animals and at 90/270 deg for the other half.

In the first experiment, we tested the orientation behavior of 18 butterflies with respect to two dark vertical stripes. All LEDs of the arena were turned blue and to generate the stripes, two sets of three LED columns were turned off (experiment: *two dark stripes)*.

Next, we recorded the headings of 18 butterflies presented with two bright stripes on a dark background. For this experiment, all LEDs were turned off and the stripes were generated by turning two sets of three LED columns blue (experiment: *two bright stripes*).

In addition, we tested 19 butterflies with respect to two artificial sun stimuli. Therefore, two LEDs at an elevation of about 23 deg were turned green (experiment: *two sun stimuli*).

### Data analysis

Heading directions were calculated by importing the data into the software MATLAB (Version R2017b, MathWorks, Natick, MA, USA) and analyzing it using the CircStat toolbox (Berens, 2009). As we changed the position of the visual stimulus every two minutes, we divided the 8-minute flights of the butterflies into four equal 2-minute sections and the 4-minute flights (stripe vs. sun stimulus) into two sections. The flight trace of each butterfly (e.g., Fig. 1B), the mean vector within each ten-second bin (Figs. 1C&F, 3B&D) and within a section (two-minute bins, Figs. 3G&I, or one-minute bins, 5A) was calculated. As the animals’ directedness can increase over the first four minutes of an experiment (Franzke et al. 2020), we focused on the butterfly’s flight performance in the last two flight sections (i.e., the last four minutes) of each experiment. Thus, the change of direction was measured as the angular difference between the mean heading directions taken during the last two flight sections (Fig. 1D, 3E&H). We then related all recorded heading angles relative to the stimulus position (stimulus position = 0 deg) and calculated the mean heading vector over the last four minutes (Fig. 2C&F, 3F, right panel). To analyze if the animals maintain a directed flight course over a shorter time period, we counted the number of ten-second bins that exceeded a directedness of *r* = 0.249 (which is the mean vector strength + 95% confidence interval for 10s bins in the last four minutes of the *no cue* experiment). For another detailed analysis of the heading distribution, we counted how often each animal kept every angle (in three deg bins) relative to the stimulus position. We defined the heading direction with the most counts as the animals’ preferred angle and normalized the number of counts of all other headings to this value. The normalized heading counts were plotted in relation to the stimulus (stimulus position = 0 deg) to generate a heatmap (e.g., 2A). For a better visualization of a bimodal or unimodal distribution of headings, we plotted the normalized heading counts in relation to the animals preferred heading (Fig. 4D). To test the butterfly’s performance in the presence of the ambiguous stimuli, we analyzed the flight trajectories over the last four minutes with a temporal resolution of two seconds. We then selected all flight sections in which the animals maintained a straight flight (<45° change in heading) over at least four seconds. We then categorized the subsequent change in flight direction according to the angular change in heading: if a butterfly changed its heading direction between 140 and 220 deg before returning to a straight flight course, this was categorized as a ‘half turn’. In contrast, a ‘full turn’ was defined when an animal changed its heading by more than 90 deg and the next phase of oriented flight deviated by less than 40° from the original direction. Based on these data, we calculated a turning index by subtracting the number of ‘full turns’ by the number of ‘half turns’ and by dividing this value by the sum of all turns. Thus, animals with a positive turning index performed more ‘full turns’ while a negative turning index represents more ‘half turns’. All butterflies that performed neither ‘full turns’ nor ‘half turns’ were excluded from this analysis.

**Fig. 2.**
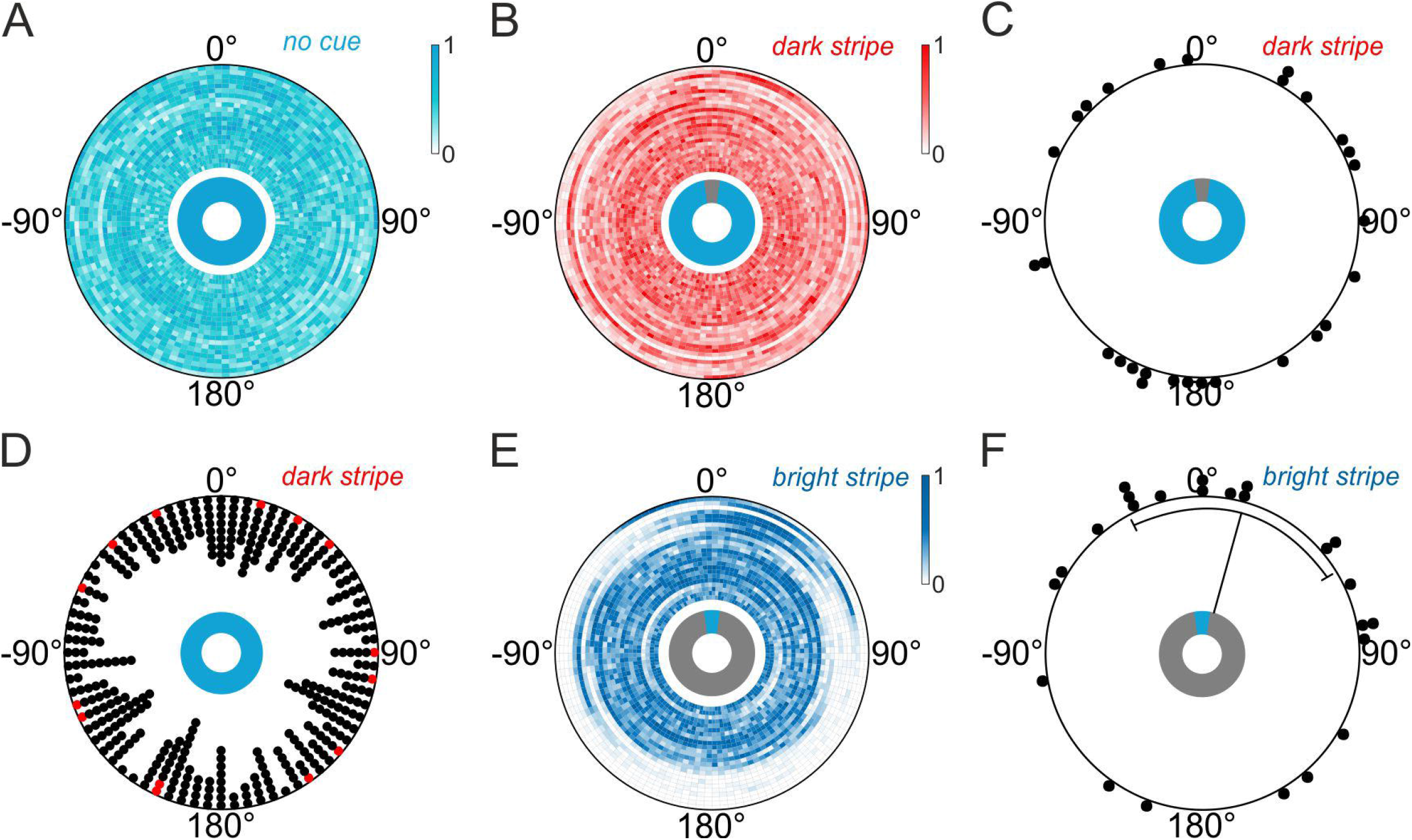
Different orientation strategies depending on the contrast of the vertical stripe. (A&B) Orientation of individual butterflies in a flight simulator either (A) without visual reference (*no cue*; N=22) or (B) with a dark stripe on a bright background (*dark stripe*; N=28) as orientation cue. The heat maps show the counted and normalized headings relative to a stimulus at 0 deg (bin size: 3 deg). Each ring represents the heading of one butterfly. The rings are sorted by increasing vector strength starting at the center. (C) Mean heading of the same butterflies relative to a dark stripe (B). Each dot in the circular plots represent the mean vector of one individual. (D) The directed phases (vector strength >0.249; bin size: 10s) in the experiment with the dark stripe are not clustered in any specific direction (*dark stripe*, middle panel). Red dots indicate the directed phases of one exemplary butterfly in arbitrary directions (P=0.774, Z=0.257, Rayleigh test). (E) Orientation of individual butterflies in a flight simulator relative to a bright stripe on a dark background (*bright stripe*; N=24). Plot conventions as in A&B. (F) Mean heading of the same butterflies relative to a bright stripe (E). Each dot in the circular plots represents the mean vector of one individual. The black lines indicate the mean and the circular standard deviation of the animals’ significant group orientation.

**Fig. 3.**
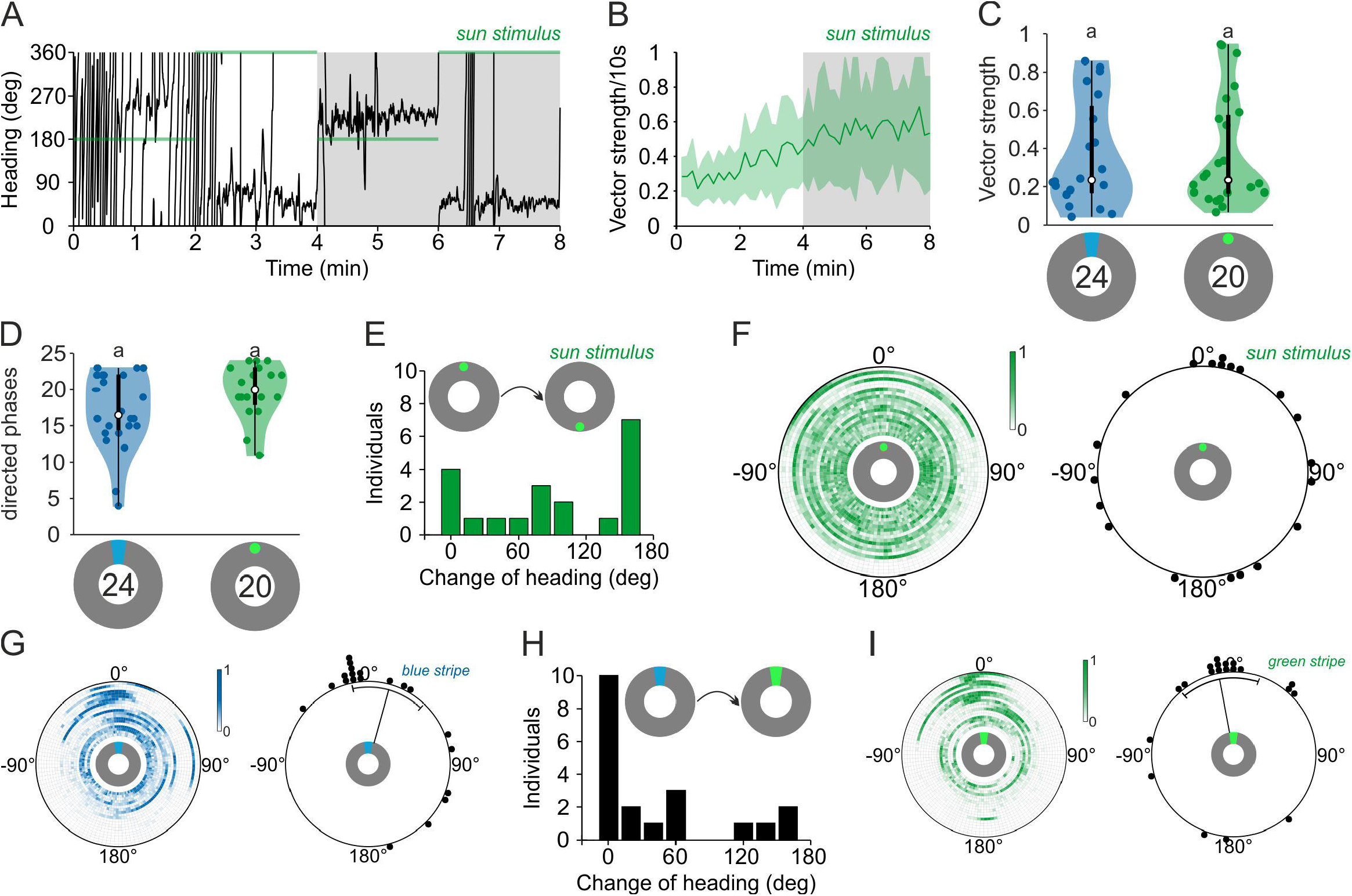
Compass orientation with respect to a sun stimulus. (A) Flight trace of one exemplary butterfly that was flying in the LED arena with respect to a simulated sun. Colored, horizontal lines indicate the position of the sun stimulus at either 0 or 180 deg. The gray boxes indicate the 4-min sections that were used for further analysis. (B) The mean vector strength *r* (bin size: 10 s) over the entire eight-minute experiments shows that the butterflies were well oriented and increased their orientation performance over time. Shaded areas indicate the 25–75% quantile. The gray boxes indicate the 4-min sections that were used for further analysis. (C) The orientation performance (bin size: 4min) did not differ when presenting either a bright stripe or a sun stimulus (P=0.759, χ^2^=0.09, Kruskal-Wallis test). White dots indicate the median vector strength. The black boxes show the interquartile range and thin black lines extend to the 1.5 x interquartile range. Colored dots show the individual data points and shaded area represents their density. Letters indicate a significant difference between the tested groups. (D) The number of oriented phases in animals flying with respect to a bright stripe was similar to the butterflies exposed to a simulated sun (P=0.110, χ^2^=2.56, Kruskal-Wallis test). Plot conventions as in C. (E) Change of heading (bin size: 20 deg) between the last two phases of an experiment (indicated by the gray boxes in A) when the stimulus was relocated by 180 deg. Most of the animals changed their heading with the displacement of the visual stimulus. (F) Orientation of butterflies (N=20) flying with respect to a simulated sun. The heat map (left panel) shows the counted and normalized headings relative to the sun at 0 deg (bin size: 3 deg). Each ring represents the heading of one butterfly. The rings are sorted by increasing vector strength starting at the center. The mean headings of the same butterflies were directed in arbitrary directions (right panel). Each dot in the circular plots represent the mean vector of one individual. (G-I) Orientation of butterflies (N=20) when the color of a bright stripe was changed from blue (G, *blue stripe*, left panels) to green (I, *green stripe*, right panels) and *vice versa*. Independent of the spectral component the butterflies flew in the direction of the bright stripe. Plot conventions as in F. The black lines in the circular plots indicate the mean and the circular standard deviation in the direction of the stripes in both experimental groups. The butterflies did not change their heading (H, bin size: 20 deg) when we changed the color of the stripe (P<0.001, u=3.047, v-test).

**Fig. 4.**
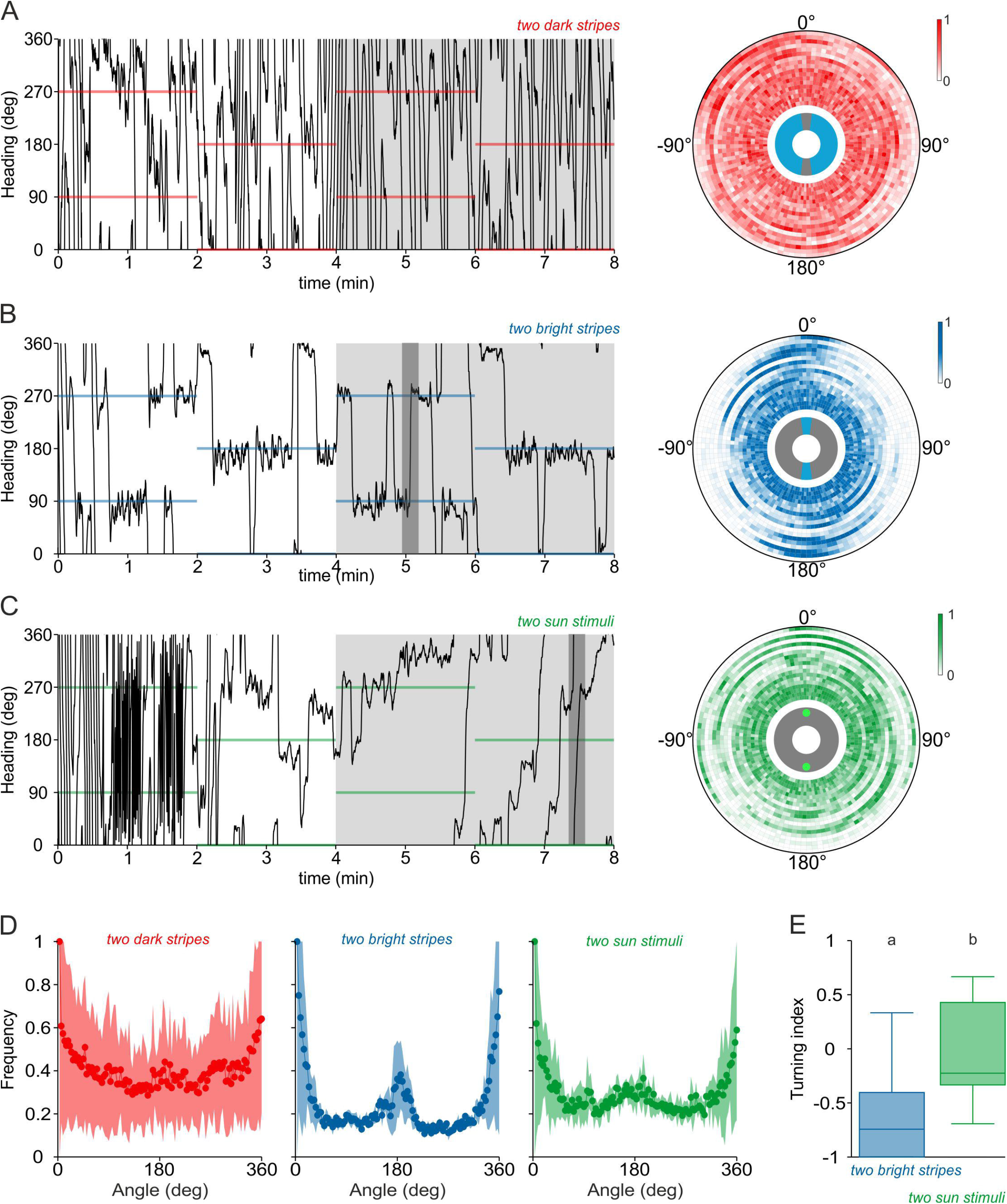
Orientation in an ambiguous scenery. (A-C) Orientation of butterflies with respect to two dark stripes on a bright background (A, *two dark stripes*; N=18), two bright stripes on a dark background (B, *two bright stripes*; N=18), or two sun stimuli (C, *two sun stimuli*; N=19). Left panels show exemplary flight trajectories of the experiments. Colored lines indicate the position of the visual stimuli at 90 and 270 deg or 0 and 180 deg. The light gray boxes indicate the 4-min sections that were used for further analysis. The dark gray part in B indicates a ‘half turn’ and the dark gray part in C indicates a ‘full turn’. Right panels: The heat maps show the counted and normalized headings relative to the stimuli shifted at 0 deg and 180 deg (bin size: 3 deg). Each ring represents the headings of one butterfly. The rings are sorted by increasing vector strength starting at the center. (D) The mean frequency of angles relative to the preferred heading of each butterfly flying with two dark stripes (first panel), two bright stripes (second panel) or two sun stimuli (third panel) as orientation reference. Butterflies perceiving two bright stripes showed a sharp second peak at 180 deg. A much wider and less high secondary peak was found at 180 deg when the animals were presented with two sun stimuli. Dots and lines represent the mean frequency and shaded areas indicate the 25–75% quantile. (E) The turning indices were calculated by dividing the number of ‘full turns’ minus the number of ‘half turns’ by the sum of all turns and differed significantly between animals tested with two bright stripes or two sun stimuli (P= 0.004, χ^2^=8.48, Kruskal-Wallis test). Horizontal lines indicate the median vector length. The boxes show the interquartile range and whiskers extend to the 2.5th and 97.5th percentiles. Letters indicate a significant difference between the tested groups.

### Statistics

The non-parametric Moore’s Modified Rayleigh test (Moore, 1980) was used to test for a bias of heading directions within a flight sector. Furthermore, we compared the heading directions of different butterfly groups with the Mardia-Watson-Wheeler test. In the experiment where we changed the color of the bright stipe, we used the v-test to test whether the butterflies kept the same heading after the stimulus manipulation. To compare the performance of the butterflies, we first calculated the mean vector strength within the last four minutes and statistically compared them using a Kruskal-Wallis-test for samples of different groups or the Wilcoxon signed-rank test for samples of the same individuals. To test whether the mean vector strength in ten-second bins increased over time, we used the Wilcoxon signed-rank test. Additionally, we compared the number of ten-second bins with a vector strength above 0.249 of different animals using a Kruskal-Wallis-Test. To test whether butterflies presented with two bright stripes, or two sun stimuli differed in their amount of ‘full turns’ vs. ‘half turns’, we compared the turning indices of both experimental groups with the Kruskal-Wallis-test.

## Results

### Landmark orientation to a vertical stripe

To investigate how monarch butterflies use a local landmark for orientation, we performed flight-simulator experiments in which individual animals were tethered at the center of an LED arena (Fig. 1A). We first presented a dark vertical stripe on a blue background to the butterflies. Although the animals were only weakly oriented, we observed sequences in which they kept a certain heading over a short time (Fig. 1B, first panel; video S1). This was different from the butterflies’ behavior in a scene without any cue (Fig. 1B, second panel). To quantify if the butterflies more often maintained constant heading directions when they had the vertical stripe as a reference, we calculated the vector strength of the mean orientation vector for each 10 s segment of the entire flight for each animal (Fig. 1C). This value ranges from 0 to 1 and indicates how directed a butterfly maintained its flight course (with 0 being completely disoriented and 1 being perfectly directed). When the butterflies had the vertical stripe for orientation, they showed a vector strength of about 0.2 ± 0.1 in the first 10 s of their flight that increased significantly to a vector strength of 0.5 ± 0.2 in the last 10 s of the flight (P=0.007, Z=-2.710, Wilcoxon signed-rank test; N=28; Fig. 1C, *dark stripe*). Without any orientation cue, the butterflies showed a vector strength of 0.2 ± 0.1 throughout the entire flight (Fig 1C, N= 22, *no cue*). This performance was significantly worse than when the vertical stripe was available as an orientation cue comparing the vector strength over the last two phases (P=0.009, χ^2^= 6.76, Kruskal-Wallis test; Fig. 1E). The higher vector strength with the vertical stripe was a result of significantly more oriented phases with *r*>0.249 (see methods; P=0.001, χ^2^= 10.31, Kruskal-Wallis test; N_*no cue*_=22, N_*dark stripe*_=28; Fig. 1F).

We also analyzed if the butterflies changed their heading when the position of the dark vertical stripe was moved by 180 deg and found that 18 out of 28 butterflies followed the stimulus relocation (directional change>90 deg; Fig. 1D, first panel). In contrast, most of the butterflies (18 of 22) tested without an orientation reference did not change their heading in a meaningful way (Fig. 1D, second panel). In summary, our data suggest that the butterflies can use a dark stripe to maintain a directed flight course.

We next tested how the butterflies use a bright stripe for orientation by inverting the visual scenery (i.e., a bright vertical stripe on a dark background). In contrast to the flight behavior with the dark stripe, many butterflies kept a constant heading over a longer time window or even over the entire 8-minute flight (Fig. 1B, third panel; video S2). This higher orientation performance was also reflected in the animals’ vector strength which significantly increased from about 0.3 ± 0.1 at the beginning to a maximum of 0.65 ± 0.3 at the end of the experiment (P<0.001, Z=-3.429, Wilcoxon signed-rank test; N=24; Fig. 1C, *bright stripe*). The vector strength of the last 4 minutes of flight was significantly higher than when the butterflies had the dark vertical stripe for orientation (P=0.004, χ^2^=8.52, Kruskal-Wallis test; Fig. 1E). Similarly, the number of oriented phases was significantly higher with the bright stripe as an orientation reference (P = 0.032, χ^2^=4.59, Kruskal-Wallis test; Fig. 1F). As expected, most of the animals (22 of 24) followed the relocation of the bright stripe (Fig. 1D, third panel) when we changed the position by 180 deg.

To gain insights into why the butterflies’ performance was different between the two experiments (bright vs. dark stripe), we next analyzed the heading directions of butterflies within the two sceneries (Fig. 2). Interestingly, animals that were tested either without a cue (Fig. 2A) or with a dark stripe (Fig. 2B) headed in all possible directions. Calculating the mean direction for each butterfly within the last four minutes relative to the dark stripe showed that the butterflies maintained arbitrary heading directions (P=0.996, R*=0.038, non-parametric Moore’s modified Rayleigh test; N_*dark stripe*_=28; Fig. 2C). However, as no butterfly maintained its direction over a longer flight sequence with the dark stripe, we studied the heading directions of the animals on a finer scale and selected only flight sequences in which the butterflies maintained a stable heading over a time window of 10 seconds. Even when we studied this, we found that the butterflies’ short-term headings were randomly distributed (P=0.774, Z=0.257, Rayleigh test; Fig. 2D), suggesting that they did not keep headings towards the stimulus. This was different from the butterflies’ behavior when a bright stripe was presented. Here, we found that most of the well-oriented animals flew in the direction of the bright stripe (P=0.003, R*=1.374, non-parametric Moore’s modified Rayleigh test; N_*bright stripe*_ =24; Fig. 2E&F), suggesting that the animals were attracted by the stimulus. Taken together, these results suggest that monarch butterflies display different behavioral strategies depending on the contrast between a vertical stripe and its background. While a dark stripe leads to several short phases of constant headings in arbitrary directions, a bright stripe allows the butterflies to maintain constant headings towards the stripe over long phases.

### Compass orientation with respect to a sun stimulus

We next wondered how monarch butterflies use a simulated sun for orientation. We therefore conducted an experiment with a green light spot as a simulated sun stimulus. Animals tested with respect to this stimulus kept constant headings over the entire experiment (Fig. 3A; video S3), although the butterflies directedness (as measured by the vector strength) significantly increased over time, from about 0.3 ± 0.1 at the beginning of the flight up to a maximum of 0.7 ± 0.3 at the end of the experiment (P=0.005, Z=-2.800, Wilcoxon signed-rank test; N=20; Fig. 3B). The vector strengths over the last four minutes of the butterflies that oriented with the sun stimulus were in the same range as the ones that had the bright vertical stripe for orientation (Kruskal-Wallis test: P=0.759, χ2=0.09; Fig. 3C). Similarly, the number of oriented phases were not significantly different between the sun-stimulus and the bright-stripe experiment (P=0.110, χ^2^=2.56, Kruskal-Wallis test; Fig. 3D). Although most of the individuals (13 of 20) changed their heading by more than 90 deg when we changed the position of the sun stimulus by 180 deg (Fig. 1E), they did not keep this stimulus in their frontal visual field. Thus, the butterflies’ heading directions were uniformly distributed (P= 0.130, R*= 0.825, non-parametric Moore’s modified Rayleigh test; N=20; Fig. 3F). This suggests that monarch butterflies can maintain any desired compass direction with respect to a sun stimulus. This difference between how butterflies treated the sun stimulus and the bright stripe was not a consequence of a difference in the spectral content (blue stripe vs. green sun): when we changed the stripe color (from green to blue and *vice versa*) every two minutes, the butterflies showed well-oriented flights, irrespective of the stripe color (Fig. 3G&I) and did not change their heading relative to the bright stripe (P=0.001, u=3.047, v-test, expectation: 0 deg; Fig 3H). In contrast, the mean direction of butterflies tested with a green stripe differed significantly from the sun-stimulus heading distribution (P=0.02, W=7.823, Mardia-Watson-Wheeler test). This indicates that the butterflies ignore the spectral content of the cue and are attracted by the brightness of the stripe while a sun stimulus is used for compass orientation.

### Orientation in an ambiguous scenery

Our previous experiments suggest that monarch butterflies may exhibit different orientation strategies depending on the appearance of a visual stimulus: they likely use a dark vertical stripe to maintain constant courses over short flight periods, while a bright stripe evokes an attraction behavior. In contrast, a simulated sun is used for a menotactic behavior, i.e., for compass orientation. Interestingly, compass orientation requires the activity of the central-complex network in fruit flies, which is not necessary for an attraction towards a stripe (Giraldo et al., 2018). To investigate in more detail whether the butterflies use different visual orientation strategies and if the central complex is likely involved in coding them, we next performed experiments within ambiguous visual scenes (two dark stripes, two bright stripes, two suns; Fig. 4). We expected that the butterflies will maintain a distinct compass heading within such ambiguous sceneries if the heading-direction network of the central complex controls the orientation behavior (Beetz et al., 2021). When we provided two dark stripes as landmarks to the butterflies, their performance resembled the performance with one dark stripe. They showed short sections of straight flights in all possible heading directions that were interrupted by rapid rotations (Fig. 4A; video S4; N=18). Again, these findings support our observation that monarch butterflies use the dark stripe/s for flight stabilization rather than for compass orientation. When the butterflies oriented with two bright stripes, they maintained a constant heading towards one of the stripes. However, they frequently switched their fixation between the stripes by changing their heading by ∼180 deg (example highlighted in dark gray in Fig. 4B, left panel; video S5). Consequently, the flight bearings were clustered around 0 and around 180 deg (Fig. 4B, right panel) which resulted in a bimodal distribution of heading directions relative to the positions of the stripes (Fig. 4D, second panel; N=18). When the butterflies were provided with the two simulated suns, they maintained arbitrary headings similar to the situation with one sun stimulus (Fig. 4C, left panel; video S6). This confirms our observation that they employ compass orientation with respect to light spots (Fig. 4C, right panel). However, we also noticed that the butterflies returned to their original bearing or headed into the opposite direction when they deviated from their course. This led to a bimodal distribution of heading directions with the second peak being less pronounced than in the two-bright-stripe experiment (Fig. 4D, third panel; N=19). To quantify if the butterflies more often returned to their original bearing when they viewed the two suns, we calculated a turning index for every butterfly. A negative turning index indicated a higher amount of 180 deg (half) turns while a positive turning index marked a higher ratio of returns to the original bearing (full turns). We found that the turning index was significantly higher with the two suns than with the two bright stripes (P= 0.004, χ^2^=8.48, Kruskal-Wallis test; N_*two bright stripes*_=17, N_*two sun stimuli*_=15; Fig. 4E). This suggests that the butterflies return to their original bearing more often when they had the two suns for orientation, a behavior that is expected if the heading-direction network of the butterfly’s central complex controls the flight direction.

### Compass orientation vs stripe attraction

As our previous experiment suggests that the butterflies’ orientation modes depended on the stimulus properties, we next wondered if the butterflies rapidly switch their orientation behavior if we changed the visual scene from a sun to a stripe stimulus (and *vice versa*). Again, when we presented a bright stripe to the butterflies, they fixated the stimulus in their frontal visual field (Fig. 5A, left panel). Interestingly, when we changed the stimulus to a simulated sun instead, the butterflies changed their heading direction and adopted arbitrary bearings with respect to the sun stimulus (Fig. 5A, right panel). The headings taken with respect to the sun stimulus were significantly different from the headings with respect to the bright stripe (P=0.002, W=12.63, Mardia-Watson-Wheeler test). Taken together, this shows that the butterflies can flexibly change their orientation strategy from compass orientation to stripe attraction during flight.

**Fig. 5.**
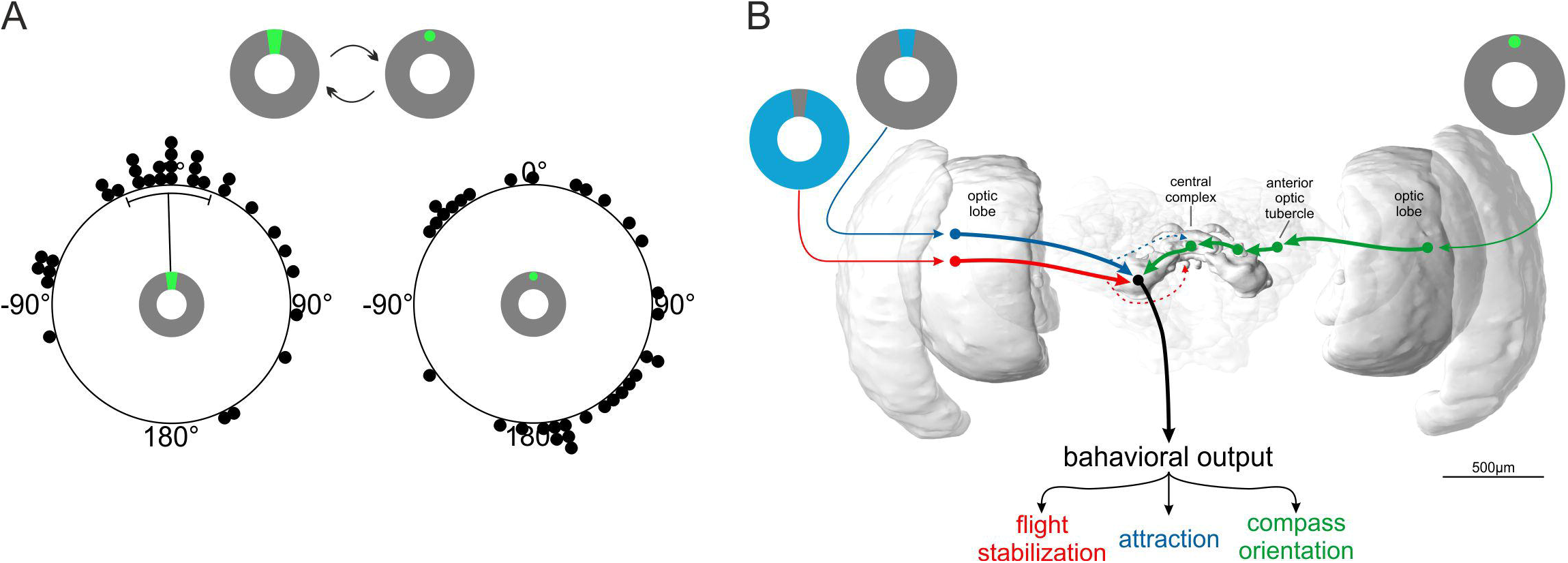
Neuronal network controlling the switch between different orientation strategies. (A) Orientation of monarch butterflies (N= 33) in a flight simulator when a bright stripe is replaced by a sun stimulus and *vice versa*. The animals switched from an attraction behavior in the direction of a bright stripe (left panel) to arbitrary directions with respect to the sun (right panel). Each dot in the circular plots represent the mean vector of one individual. The black lines indicate the mean and the circular standard deviation of the animals’ significant group orientation. (B) Schematic illustration of the proposed different neuronal pathways resulting in different orientation strategies. While visual information about landmark cues (red and blue pathway) is directly transferred from the optic lobes to the lateral accessory lobes, information of the position of the sun (green pathway) is first integrated in the animals’ internal compass system, the central complex, by passing the anterior optic tubercle and the bulb. Compass information from the central complex is sent to the lateral accessory lobe. From there, behavioral output is driven via descending neurons (black arrow) resulting in different orientation strategies depending on the visual input. The brain of the monarch butterfly is adapted from Heinze and Reppert (2012) and created via insectbraindb.org (Heinze et al., 2021).

## Discussion

We here tested the ability of monarch butterflies to use different visual stimuli to maintain a directed flight course and found that they exhibit different orientation modes that depend on the stimulus identity. Vertical stripes led to a rather simple strategy based on flight control or stripe fixation while a simulated sun evoked compass orientation. This suggests that different strategies operate in parallel in the brain (Fig. 5B) which allows monarch butterflies to effectively adapt their orientation strategy to a certain behavior by dynamically switching to the most appropriate strategy during flight.

### Orientation to local cues

A bright stripe triggered an attraction behavior in monarch butterflies. We interpret this behavior as a brightness-based flight approach with the intention to leave the current setting and access a new environment similar to what has been found in navigating orchid bees (Baird and Dacke, 2016). This would also be in line with our observation that the behavior does not seem to be affected by the stripe’s spectral information. However, instead of centering the bright stripe accurately in their frontal visual field, many butterflies kept the stripe slightly to their left/right vertical body axis. This indicates that the butterflies rely on the edge between the stripe and the background to sustain a constant heading, similar to what has been shown in walking *Lucilia* flies (Osorio et al., 1990). The stripe fixation of the butterflies that we described here is well in line with the results reported for tethered-flying *Drosophila* that are also attracted by a bright stripe (Giraldo et al., 2018; Maimon et al., 2008). However, the fly’s positive taxis seem to be dependent on the behavioral or locomotory state. Thus, walking fruit flies can also adopt arbitrary headings with respect to a bright stripe (Green et al., 2019). Interestingly, in flying and walking fruit flies (Götz, 1987; Horn and Wehner, 1975; Strauss and Pichler, 1998) and other insects, such as flying locusts (Baker, 1979; Robert, 1988) or naïve walking ants (Buehlmann et al., 2020), a dark stripe also elicits stripe fixation. In contrast, monarch butterflies used the dark stripe to occasionally maintain a bearing over short phases in a random direction. This result is similar to what has been reported for monarch butterflies in a more complex visual scene, where they had the panoramic skyline for orientation (Franzke et al., 2020). Interestingly, in the same study the butterflies showed a similar behavior when they experienced a grating pattern – providing rotational optic flow – as the only visual input in a flight simulator (Franzke et al., 2020). Such rotational optic flow can provide an animal with directional information relative to a visual cue to perform compensatory steering and to keep a certain bearing (Wolf and Heisenberg, 1990; Zeil, 1996; Zeil et al., 2008). Thus, instead of using the vertical dark stripe to maintain a desired heading over an entire flight, our data suggest that the butterflies use optic-flow information to stabilize their heading over short flight sequences. In summary, a dark stripe evokes a different behavior in monarch butterflies than a bright stripe, which stands in contrast to *Drosophila*. It will be interesting to observe in the future at which visual angle of the bright stripe the butterflies will switch to an attraction behavior, and at which stripe width flight control will dominate the orientation behavior in monarch butterflies.

### Sun compass orientation

When we presented a simulated sun to the butterflies, they kept arbitrary headings relative to the stimulus. This menotactic behavior is in line with what has been reported for other insects such as fruit flies (Giraldo et al., 2018) or dung beetles (Byrne et al., 2003). In theory, menotaxis can be carried out by a simple, vision-based retina matching of the current and remembered sun position similar to how many insects can use the profile of a panoramic skyline for orientation (Cartwright and Collett, 1983; Collett, 1992; Junger, 1991; Lent et al., 2010; Wehner and Räber, 1979). However, when we provided the butterflies with two simulated suns set 180 deg apart as orientation references, they returned to their original bearing during flight, which shows that they compute a distinct heading direction with respect to the ambiguous visual scene. This observation suggests that they do not only rely on the azimuth of the sun stimulus for orientation but their orientation mechanism requires the involvement of the activity of a multisensory heading-direction network. This raises the question of what exactly defines the green light spot as a compass cue. In a recent paper, the butterflies’ flight headings were directed towards the sun stimulus when the elevation of the sun stimulus was set to a low elevation of about 5 deg (Franzke et al., 2020). Even though the contrast between the background and the sun stimulus in Franzke et al. (2020) might have led to these heading choices, it opens up the possibility that the elevation of the sun stimulus is a critical parameter to induce compass orientation. In addition, for maintaining a certain heading direction, compass orientation also requires the network to memorize the desired direction (Grob et al., 2021; Honkanen et al., 2019). Whether the monarch butterfly can develop a long-term memory for a direction relative to the sun stimulus, as shown in the fruit fly (Giraldo et al., 2018) awaits to be investigated. Similarly, our future studies will focus on the use of the sun stimulus in the context of migration. Rather than adopting arbitrary headings, we expect that migratory monarch butterflies keep directed courses with respect to the sun stimulus that would guide them to the migration destination. Moreover, as the butterflies employ a time-compensated sun compass during their migration (Merlin et al., 2009; Mouritsen and Frost, 2002), we will next study if the heading to the sun stimulus will be adjusted according to the time-of-day. Taken together our findings show, that monarch butterflies use a sun stimulus for compass orientation, a strategy that allows them to maintain any arbitrary heading with respect to the sun during dispersal or in a distinct southward direction when they are in their migratory stage.

### Neuronal network behind orientation

Our experiments suggest that the butterfly brain generates different orientation strategies but how is this accomplished at the neuronal level? As the butterflies used the dark stripe for flight control, the neuronal basis for it lies likely in the motion vision center, the lobula plate of the optic lobe (Meier and Borst, 2019; Ullrich et al., 2015). Although some optic-flow information is integrated into the central complex in locusts and bees (Rosner et al., 2019; Stone et al., 2017), the relevant information for flight control is directly transferred to the thoracic ganglia via descending pathways (Suver et al., 2016). In fruit flies, attraction does not require the activity of the central complex (Giraldo et al., 2018). This is well in line with our results from the 2-bright stripe experiment which points towards a coding of directional information without the association of a multisensory heading-direction network in monarch butterflies. Thus, the basis for the attraction to a bright stripe might also be based on the motion-vision network that is directly connected to descending neurons, as suggested in a recent model (Fenk et al., 2014) (Fig. 5B). In contrast, the butterflies resolved the ambiguity of the visual scene, when we instead presented two suns as stimuli for orientation. This matches recordings from the heading-direction network in the butterfly central complex that encodes an explicit heading based on multisensory inputs if confronted with a similar 2-sun stimulus (Beetz et al., 2021). Thus, our behavioral data suggest that the central complex encodes the sun stimulus, which is also in line with the sensitivity of central complex neurons to a green light spot (Heinze and Reppert, 2011; Nguyen et al., 2021). Recent results suggest that the central complex compares the actual heading direction with the desired direction (Green et al., 2019; Stone et al., 2017). By encoding the desired migratory direction, the butterfly’s central complex is likely taking a central role in the migration and is the region in the brain where time-of-day information becomes relevant for sustaining the migratory southward direction. We therefore propose that compass orientation is processed by the central complex, while stripe attraction and flight control seem to rely on reflexive pathways without the involvement of a higher brain center (Fig. 5B). Our results here show that the butterflies can switch between compass orientation and attraction. Information from the central complex is sent to the lateral accessory lobe and further to the posterior protocerebrum in monarch butterflies (Heinze et al., 2013) where it might converge with the attraction and flight control pathways (Fig. 5B). Interestingly, recent results in the fruit fly suggest that descending neurons can generate different steering commands based on different input pathways (Rayshubskiy et al., 2020). This suggests that the reliance on different orientation strategies might also be weighted and governed by descending neurons in monarch butterflies, which allows butterflies to rapidly switch their orientation strategy during flight. This enables them to flexibly switch from a long-distance system during dispersal or migration to a short-distance orientation strategy such as the attraction to their host plant.

## Author contributions

Study design: MF, KP, BeJ. Conducting experiments: MF, MG, CK. Analysis of data: MF, DD, BeJ. Interpretation of data: MF, KP, BeJ. Drafting of the manuscript: MF, BeJ. Critical review of the manuscript: CK, MG, DD, KP. Acquired Funding: BeJ. All authors approved of the final version of the manuscript.

## Competing interests

The authors declare no competing interests.

## Funding

This work was supported by the Emmy Noether program of the Deutsche Forschungsgemeinschaft granted to BeJ (GZ: EL784/1-1).

## Acknowledgments

We thank Jerome Beetz, James Foster, and Anna Stöckl for fruitful discussions on the manuscript. We are grateful to Konrad Öchsner for his help in designing the LED arena as well as the LED band for simulating the sun. We also thank the mechanics workshop of the Biocenter (University of Wuerzburg) for building important pieces of the flight simulator. In addition, we would like to thank Sergio Siles (butterflyfarm.co.cr) and Marie Gerlinde Blaese for providing us with monarch butterfly pupae.

## Figure Legends

**Table 1. Properties of the different presented stimuli**. The table summarizes the proportion, color, and intensity of each stimulus as well as the color and intensity of the background of the LED arena.

**Video S1**. Demonstration of a butterfly tethered at the center of a flight simulator and flying with respect to a dark stripe on a bright background. The position of the stimulus was relocated by 180 deg every two minutes.

**Video S2**. Demonstration of a butterfly tethered at the center of a flight simulator and flying with respect to a bright stripe on a dark background. The position of the stimulus was relocated by 180 deg every two minutes.

**Video S3**. Demonstration of a butterfly tethered at the center of a flight simulator and flying with respect to a bright green light spot as sun stimulus. The position of the stimulus was relocated by 180 deg every two minutes.

**Video S4**. Demonstration of a butterfly tethered at the center of a flight simulator and flying in an ambiguous scene with two dark stripes on a bright background. The position of the stimuli was relocated by 90 deg every two minutes.

**Video S5**. Demonstration of a butterfly tethered at the center of a flight simulator and flying in an ambiguous scene with two bright stripes on a dark background. The position of the stimuli was relocated by 90 deg every two minutes.

**Video S6**. Demonstration of a butterfly tethered at the center of a flight simulator and flying in an ambiguous scene with two bright green light spots as sun stimuli. The position of the stimuli was relocated by 90 deg every two minutes.

## Notes

### Competing Interest Statement

The authors have declared no competing interest.

